# E3 ubiquitin ligase SYVN1 mediates K63-linked ubiquitination of DDX3X to activate Macrophage NLRP3 Inflammasome

**DOI:** 10.64898/2026.09.15.751797

**Authors:** Mohammad Anas, Abhalaxmi Singh, Nithish Raj Prasad, Joshua W. Thompson, Chinnaswamy Tiruppathi, Asrar B. Malik

**Affiliations:** Department of Pharmacology and the Center for Lung and Vascular Biology, The University of Illinois College of Medicine, Chicago, IL, 60612, USA; Cell Biologics, 2201 W Campbell Park Drive, Chicago IL, 60612

## Abstract

DDX3X (DEAD-box helicase 3, X-linked) is a common and essential component for both stress granules and NLRP3 inflammasome assembly and their activation; however, the upstream cellular stress signals driving DDX3X to activate the contrasting cellular pathway remain unclear. We identified the pivotal role of the E3 ubiquitin ligase SYVN1(Synoviolin) as the upstream regulator of DDX3X and thereby controlling NLRP3 activation and stress granule assembly. We observed SYVN1 silencing in macrophages prevented both NLRP3-driven inflammation and stress granule formation. SYVN1 deficiency prevented LPS-induced inflammatory lung injury and increased the survival rate of the mice. SYVN1 sustains DDX3X gene expression and promotes stimulus-dependent ubiquitination of DDX3X. Under inflammatory conditions, SYVN1 mediated 63-linked ubiquitination of DDX3X, a requirement for NLRP3 inflammasome activation. Conversely, stress conditions reduced K63-linked DDX3X ubiquitination in coordination with activity of the deubiquitinase OTUB1 (OTU domain-containing ubiquitin aldehyde-binding protein 1). Thus, the balance between SYVN1 and OTUB1 functioned to optimize DDX3X activity and activation of NLRP3 or stress granule. These findings show the upstream role of SYVN1–OTUB1 axis in integrating cellular stress signals to decide the cell fate and suggest that ubiquitination of DDX3X is a potential target for inflammasome driven inflammation.

## Introduction

Inflammasomes are cytosolic multi-protein complexes that serve as critical sensors of pathogen-associated molecular patterns (PAMPs) and danger-associated molecular patterns (DAMPs), coordinating innate immune responses through caspase-1 activation and downstream cytokine maturation [1–4]. Among the best-characterized inflammasomes, the NLRP3 (NOD-like receptor pyrin domain-containing 3) inflammasome plays a central role in macrophage-mediated inflammatory responses [3]. NLRP3 inflammasome activation requires two sequential steps: a priming phase characterized with NF-KB–dependent transcriptional upregulation of NLRP3, pro-IL-1β, and pro-IL-18 through TLRs (Toll like receptors); and an activation phase, triggered by potassium efflux–dependent signals (LPS with nigericin or ATP) or by potassium efflux– independent signals such as imiquimod that directly bind with TLR7 and induce the inflammasome [3, 5–7]. Activated caspase-1 processes pro-IL-1β and pro-IL-18 into their mature, secreted forms and cleaves gasdermin D (GSDMD) releasing N-terminal fragments oligomerize to form lytic pores in the plasma membrane that lead to pyroptosis [8, 9]. Although NLRP3 inflammasome activation is essential for host defense against infection, its dysregulation contributes to a wide range of inflammatory diseases, including sepsis, acute lung injury, atherosclerosis, type 2 diabetes, and Alzheimer’s disease[3, 4, 10, 11]. While the NLRP3 inflammasome serves as a critical complex for innate immune signaling, cells also utlizes distinct adaptive mechanism to respond to diverse cellular stress including oxidative stress, heat shock, viral infection, hypoxia, and DNA damage can rapidly assemble cytoplasmic ribonucleoprotein complexes known as stress granules (SGs) [12, 13]. Stress granules are membrane less organelles formed through liquid– liquid phase separation (LLPS), comprising stalled 48S preinitiation complexes, RNA-binding proteins, and associated mRNAs[12, 14]. Stress granules are broadly considered as a pro-survival response that allows cells to redirect resources toward stress adaptation [13]. Upon resolution of the stressor (e.g., heat shock), stress granules disassemble through K63-linked polyubiquitination of G3BP1 (Ras GTPase-activating protein-binding protein 1) by the ER-associated adaptor FAF2 and the ubiquitin-dependent AAA+ ATPase segregase VCP/p97 [15]. Dysregulation of stress granule assembly and disassembly dynamics has been linked to several neurological diseases, including amyotrophic lateral sclerosis (ALS), Huntington’s disease, and cancer progression [16–18].

Emerging evidence indicates that the NLRP3 inflammasome and stress granules are functionally interconnected and compete for shared molecular machinery [21]. A central node in this crosstalk is DDX3X, an ATP-dependent RNA helicase that acts as a critical component and is required for activation of the NLRP3 inflammasome or stress granules [21]. DDX3X directly interacts with NLRP3 to promote inflammasome activation and ASC speck formation, whereas during non-inflammatory stress, DDX3X sequesters to assemble stress granules [21]. These findings highlight the importance of DDX3X in maintaining the cellular homeostasis [17]; however, the critical unaddressed question is how DDX3X interprets these signals to determine cell fate in response to various stress stimulus.

Evidence highlights the importance of ubiquitination as a key regulatory step governing diverse cellular signaling pathways, including innate immune and distinct stress related responses such as assembly, disassembly and activation of NLRP3 inflammasome and stress granules [22–25]. E3 ubiquitin ligases confer specificity and distinct functional outcomes by directing attachment of ubiquitin chains to their target substrates. Among the different types of ubiquitin chains, the role of K48 and K63-linked ubiquitin chains are most predominant in regulating and function of target proteins. K48 linked ubiquitin chains target proteins for proteasomal degradation whereas K63-linked chains induce non-degradative functions including signal transduction, suppression of protein and protein–protein interactions [26–29]. Importantly, K27-linked chains are also implicated in innate immune signaling and DNA damage responses [26, 30, 31]. Among E3 ligases involved in regulating inflammatory and stress responses, SYVN1 (synoviolin 1), also known as HRD1, has emerged as a multifunctional regulator of cellular homeostasis [40]. SYVN1 is a RING-domain E3 ubiquitin ligase canonically associated with endoplasmic reticulum-associated degradation (ERAD), a quality-control pathway that retrotranslocates misfolded ER proteins for ubiquitin–proteasome-mediated degradation [32]. SYVN1 was originally identified as overexpressed in synovial fibroblasts of rheumatoid arthritis patients, where it promoted joint inflammation and prevented apoptosis of hyperplastic synoviocytes [33–35]. Studies also show that SYVN1 targets diverse substrates, proapoptotic factor IRE1 and B lymphocyte-induced maturation protein 1 (BLIMP1) establishing it in immune homeostasis [34, 36]. SYVN1 additionally regulates TLR4-induced inflammation through ubiquitination and inactivation of USP15, thus amplifying NF-KB-mediated inflammatory gene expression [37]. A recent study also identified SYVN1 as a direct E3 ubiquitin ligase for GSDMD that promotes K27-linked ubiquitination of GSDMD at residues K203 and K204 and facilitates pyroptotic cell death [38]. Although SYVN1 is implicated in regulating distinct cellular signaling pathways mediating inflammation, it is unknown whether it controls the activation of the NLRP3-stress granule node. Such a mechanism could enable SYVN1 as check point molecular integrator that interprets signals to determine cell fate.

In this study, we identify SYVN1 as a key regulator of NLRP3 inflammasome activation and stress granule assembly through ubiquitination dependent activation of DDX3X. In addition to ubiquitin ligases, deubiquitinases (DUBs) provide an additional layer of regulation by counterbalancing E3 ubiquitin ligases through the removal of the ubiquitin chains from the targeted proteins. Consistently, several DUBs have been shown to regulate NLRP3 inflammasome activation and stress granule dynamics such as BRCC3, USP7, USP47, USP13 and USP1/UAF1 regulates NLRP3 inflammasome assembly and USP5 and USP13 regulates stress granule assembly [43–47]. We show that the deubiquitinase OTUB1 to be a critical regulator of the SYVN1-DDX3X axis, demonstrating that OTUB1 cooperates with SYVN1 to modulate DDX3X ubiquitination and tune cell fate in response to stress stimulus. Together, our findings identify a SYVN1-OTUB1 regulatory axis that integrates ubiquitin dependent signaling to determine cellular fate and maintain homeostasis. This study also identifies SYVN1 as therapeutic target with potential implications for inflammatory diseases such as sepsis and acute lung injury.

## Results

### SYVN1 is required for the activation of NLRP3 Inflammasome

LPS stimulates the expression of SYVN1 in bone marrow derived macrophages (BMDMs) indicating the importance of SYVN1 in inflammation (Figure 1a). To understand the role of SYVN1 in inflammation, BMDMs were primed with LPS and stimulated with the canonical NLRP3 activators nigericin or ATP. Notably, SYVN1 depletion significantly reduced the secretion of IL-1β and IL-18, indicating that SYVN1 is required for NLRP3 inflammasome activation (Figure 1b–c). Furthermore, SYVN1 deficiency impaired ASC oligomerization which was corroborated by glycerol gradient fractionation showing decreased ASC oligomerization in high-molecular-weight fractions (Figure 1d-e). Consistent with these results, both ASC speck formation and the percentage of speck-positive cells were significantly reduced in SYVN1-deficient BMDMs (Figure 1g). Together, these findings demonstrate that SYVN1 is required for NLRP3 inflammasome activation.

**Figure 1.**
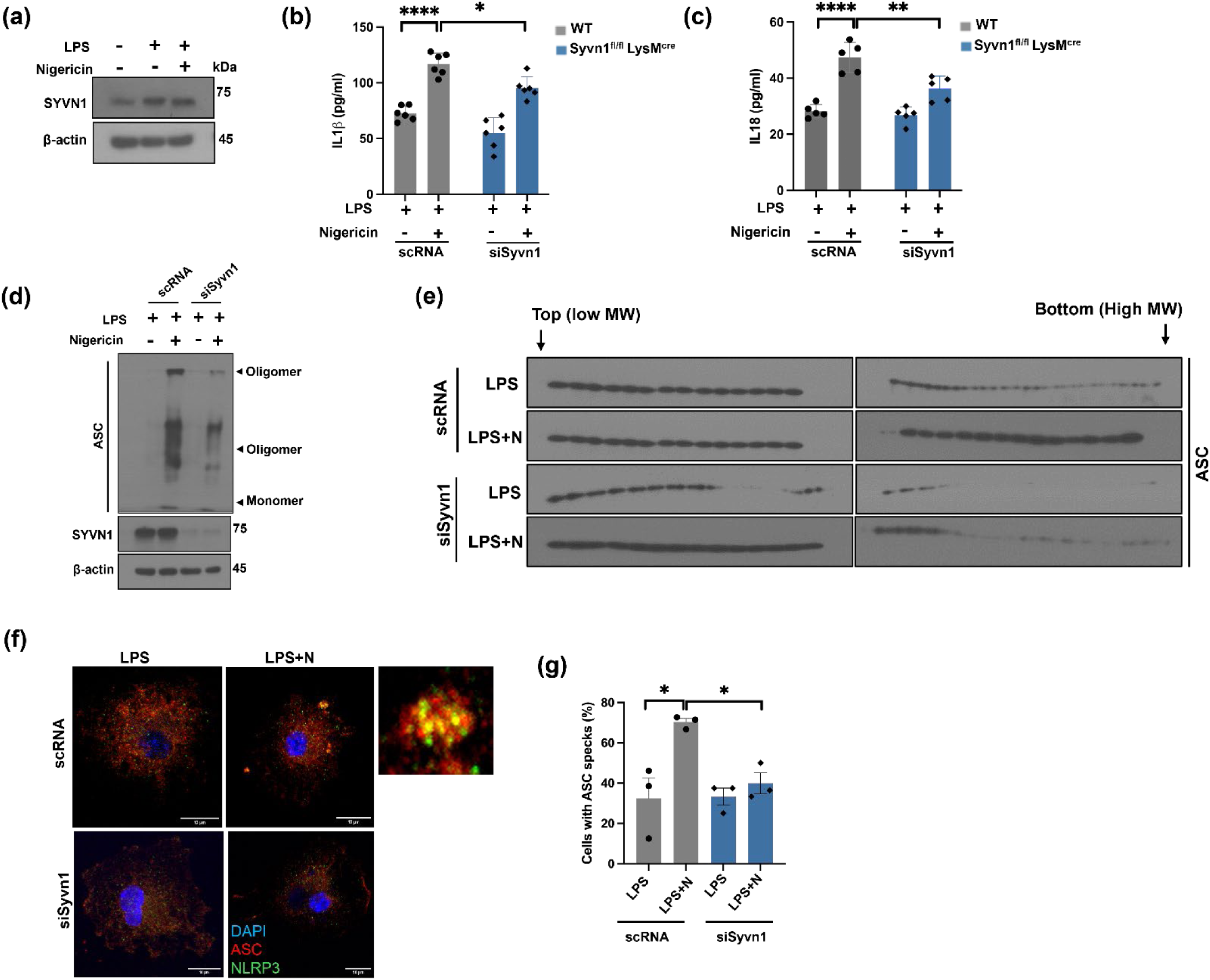
Depletion of SYVN1 prevents NLRP3 inflammasome activation in macrophages. (a) Immunoblot analysis of SYVN1 expression after LPS stimulation (representative blots, n=3). (b-c) Enzyme-linked immunosorbent assay measurement of IL-1β and IL-18. P values: *p<0.05, **p<0.01 and ****p<0.0001. (d) Immunoblot analysis of ASC oligomerization from the insoluble fraction in non-reducing conditions (representative blots, n=3). (e) Immunoblot analysis of lysates fractionated in 15-45% glycerol gradient by analytical ultracentrifugation to assess the oligomerization of ASC in BMDMs stimulated with 1μg/ml LPS (3h) with or without 20μM nigericin (30 min) in control and SYVN1 depleted BMDMs (representative blots, n=2). (f-g) Confocal images of NLRP3 inflammasome assembly after siRNA mediated knockdown of Syvn1 in BMDMs. Scale bars,10μm (whole cell images); 1μm (magnified images) (representative images, n=3). Quantification of ASC speck positive cells stimulated with LPS and nigericin as indicated. P value: *p<0.05. Data are mean ± s.e.m.

### SYVN1 interacts with DDX3X and controls its expression transcriptionally

To identify SYVN1-interacting proteins relevant to NLRP3 inflammasome regulation, we performed affinity purification coupled with mass spectrometry (AP-MS) using Flag-tagged SYVN1 expressed in HEK293T cells (Figure 2a). This approach identified DDX3X, an RNA helicase known to function as a component of both the NLRP3 inflammasome and stress granules. Co-immunoprecipitation confirmed the SYVN1–DDX3X interaction (Figure 2b). Notably, among canonical inflammasome components, only NLRP3 associated with SYVN1 (Figure 2b). Moreover, SYVN1 did not interact with G3BP1, a canonical stress granule marker as well indicating the specific interaction with DDX3X (Figure 2b). We next mapped the domains of both SYVN1 and DDX3X responsible for the interaction. SYVN1 full length and N-terminal containing TM and RING domain interacted with DDX3X. Reciprocally, all DDX3X domains interacted with SYVN1, suggesting a broad distributed binding surface (Supplementary figure 4a-d). Since DDX3X is a limiting, shared component, its availability determines the activation of NLRP3 inflammasome or stress granule depending on the stimulus [18]. We hypothesized whether SYVN1 regulates both NLRP3 inflammasome and stress granule assembly by modulating DDX3X in macrophages. To test that, we generated myeloid-specific SYVN1-knockout mice by crossing Syvn1^fl/fl^ with LysM-cre mice (Supplementary figure. 1a-b). Syvn1^fl/fl^LysM^cre^ mice displayed no apparent defects in hematopoietic development (Supplementary figure 1c). Strikingly, loss of SYVN1 markedly reduced DDX3X protein abundance in macrophages (Figure 2c), a phenotype recapitulated by siRNA-mediated SYVN1 depletion (Supplementary figure 2a). In contrast, depletion of DDX3X did not alter SYVN1 expression (Supplementary figure 2b), supporting a unidirectional regulatory relationship. We next asked whether SYVN1 controls DDX3X expression at the transcriptional or post-translational level. SYVN1 depletion significantly reduced Ddx3x mRNA, whereas DDX3X depletion had no effect on Syvn1 gene expression (Figure 2d; Supplementary figure 2c). Moreover, cycloheximide-chase analysis revealed that DDX3X abundance was already reduced at the onset of the assay in SYVN1-deficient cells, without evidence of accelerated protein decay (Figure 2e). Consistent with this, pharmacological inhibition of proteasomal degradation with MG132 failed to restore DDX3X protein levels (Figure 2f). Together, these data indicate that SYVN1 sustains DDX3X abundance primarily by promoting its gene expression rather than by stabilizing the DDX3X protein. Thus, SYVN1 emerges as an upstream regulator of the DDX3X to regulate NLRP3 inflammasome and stress-granule responses.

**Figure 2.**
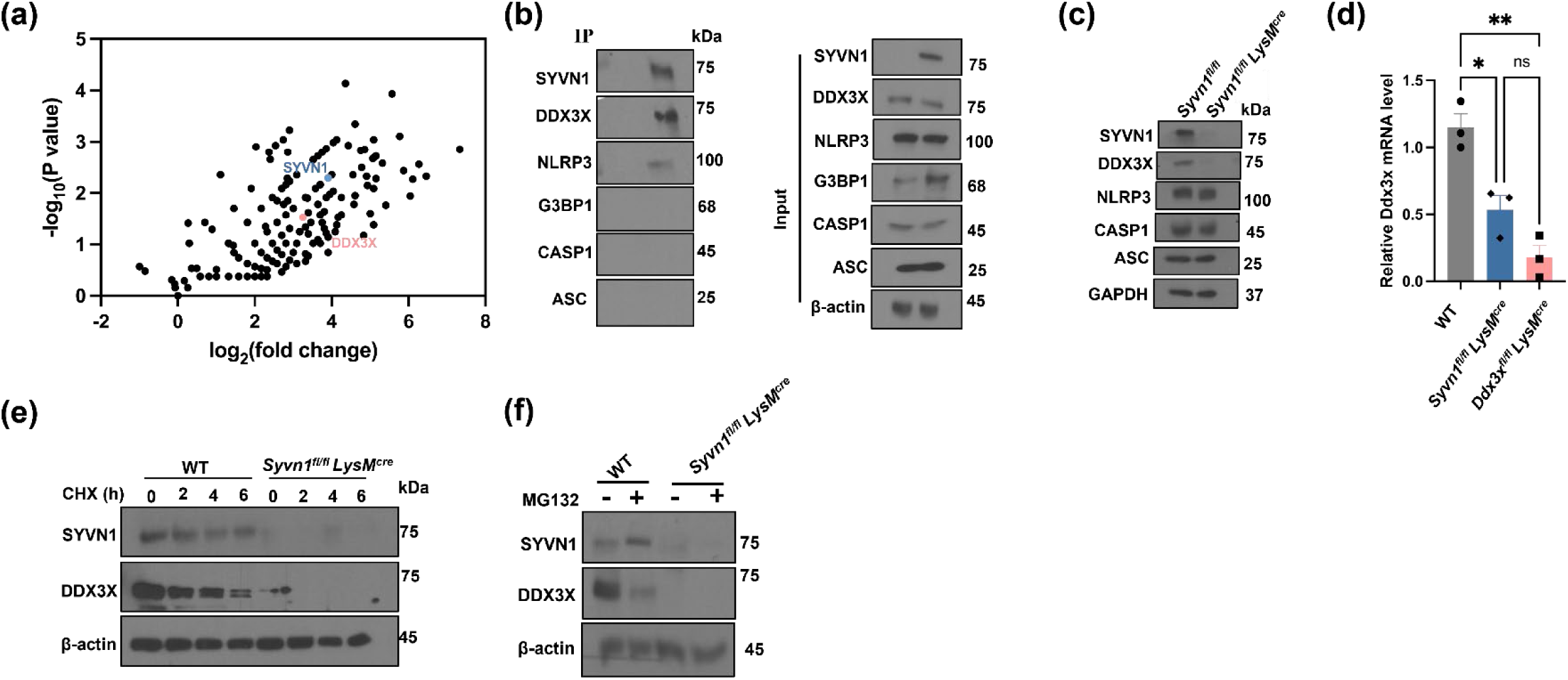
SYVN1 directly interacts with DDX3X and controls DDX3X protein level transcriptionally. (a) Volcano plot from mass spectrometry analysis of proteins co-immunoprecipitated with SYVN1. SYVN1 (steel blue) and DDX3X (salmon pink) are highlighted (n=2). (b) Co-immunoprecipitation of SYVN1 and DDX3X in BMDM treated with LPS (representative blots, n=3). (c) Immunoblot analysis of SYVN1, DDX3X, NLRP3, Caspase1, ASC expression in wild type (WT) and SYVN1 depleted macrophage cells (representative blots, n=3). (d) qPCR analysis of the mRNA levels of DDX3X in WT and Syvn1^fl/fl^LysM^cre^ (n=3). P values: *p<0.05, **p<0.01 and ns = not significant. (e) Immunoblot analysis of SYVN1 and DDX3X expression in wild type (WT) and SYVN1 depleted macrophage cells after treatment with 100μg/ml cycloheximide at the indicated time points. (representative blots, n=3). (f) Immunoblot analysis of SYVN1 and DDX3X expression in wild type (WT) and SYVN1 depleted macrophage cells after treatment with 10μg/ml MG132 for 2 h (representative blots, n=3).

### SYVN1 deficiency impairs inflammasome activation and protects against LPS-induced lung injury

Functionally, SYVN1 deficiency markedly impaired NLRP3 inflammasome activation in BMDMs, as evidenced by reduction in caspase-1 activation and GSDMD cleavage, IL-1β and IL-18 secretion following inflammasome stimulation (Figure 3a, Supplementary figure 3e-f). This defect was observed across diverse NLRP3 agonists, including LPS combined with ATP, imiquimod, and Pam3CSK4 or poly(I:C) combined with nigericin (Supplementary figure 3a-d) indicating that SYVN1 acts broadly within the NLRP3 pathway rather than at the level of a specific upstream stimulus. Consistent with this, both SYVN1- and DDX3X-deficient BMDMs exhibited impaired ASC oligomerization (Figure 3b) placing SYVN1–DDX3X regulation upstream of inflammasome assembly. SYVN1 deficiency also markedly reduced ASC speck formation and stress-granule assembly attributed to the loss of DDX3X (Supplementary figure 3g-h).

**Figure 3.**
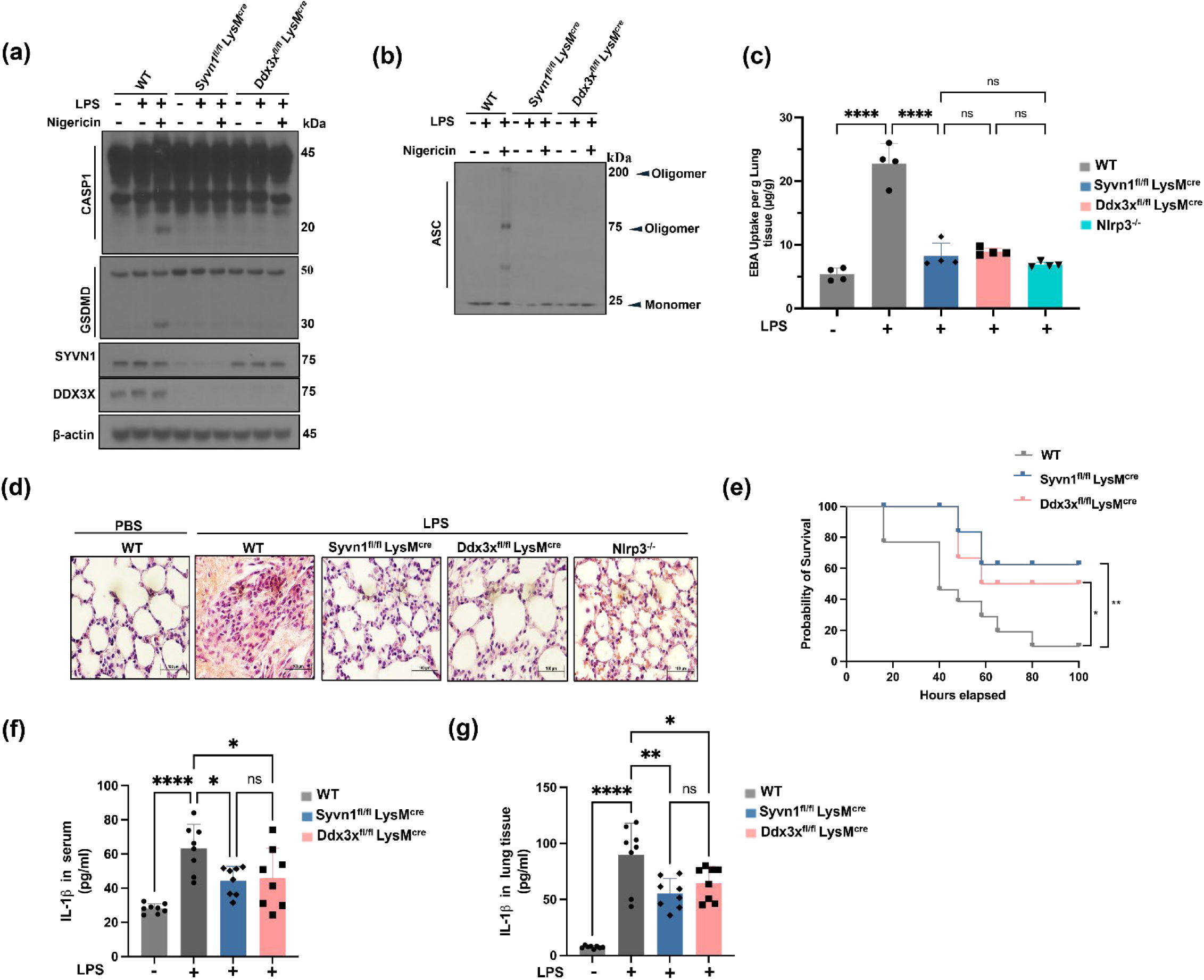
SYVN1 promotes NLRP3 inflammasome activation through DDX3X. (a) Immunoblot analysis of CASP1 cleavage and GSDMD cleavage in WT, Syvn1^fl/fl^ LysM^cre^, Ddx3x^fl/fl^ LysM^cre^ BMDMs after treatment of LPS with or without nigericin (representative blots, n=3). (b) Immunoblot analysis of cross-linked ASC oligomers in WT, Syvn1^fl/fl^ LysM^cre^, Ddx3x^fl/fl^ LysM^cre^ BMDMs after treatment with LPS with or without nigericin (representative blots, n=3). (c) Lung vascular permeability to measure the injury (EBA uptake in lungs) was determined in Syvn1^fl/fl^ LysM^cre^, Ddx3x^fl/fl^ LysM^cre^ and Nlrp3^−/-^ and compared with WT (n=4). P values: ****p<0.0001 and ns = not significant (d) Images of inflammatory cells in alveoli per 4mm^2^ (using the Fiji image analysis) in WT, Syvn1^fl/fl^ LysM^cre^ and Ddx3x^fl/fl^ LysM^cre^ lung after injecting LPS to evaluate lung injury (representative images, n=5). (e) Assessed survival of the mice after LPS (15mg/kg) injection peritoneally in WT, Syvn1^fl/fl^ LysM^cre^ and Ddx3x^fl/fl^ LysM^cre^ mice (n=8). P values: *p<0.05 and **p<0.01. (f-g) Enzyme-linked immunosorbent assay measurement of IL-1β in serum and lung tissue from WT, Syvn1^fl/fl^ LysM^cre^ and Ddx3x^fl/fl^ LysM^cre^ mice (n=8). P values: *p<0.05, **p<0.01 and ****p<0.0001.

We next examined the physiological relevance of SYVN1-dependent inflammasome regulation in vivo. Following intraperitoneal LPS challenge, SYVN1 and DDX3X deficient mice were protected from LPS-induced pulmonary injury and displayed significantly improved survival (Figure 3 c-e). Consistent with attenuated inflammasome activation, myeloid-specific SYVN1 and DDX3X-deficient mice exhibited substantially lower IL-1β levels in both serum and lung tissue than control mice (Figure 3 f-g). Together, these findings establish SYVN1 as a critical upstream regulator of DDX3X-dependent NLRP3 inflammasome and stress granule assembly and their activation.

### SYVN1 mediated K63-linked ubiquitination of DDX3X directs the NLRP3 inflammasome–stress granule switch

Mass spec and co-immunoprecipitation data showed strong interaction between SYVN1 and DDX3X indicating SYVN1 not only controled the gene expression of DDX3X but it interacted with DDX3X protein and modulates its activity (Figure 2a-b). To test this hypothesis, HEK293T cells were co-transfected with SYVN1-Flag, DDX3X-His, and HA-tagged ubiquitin, followed by co-immunoprecipitation. SYVN1 promotes ubiquitination of DDX3X suggesting posttranslational regulation of DDX3X by SYVN1 (Figure 4a). To delineate the mechanism, we defined the specific linkage of ubiquitination on DDX3X. HEK293T cells were co-transfected with SYVN1-Flag, DDX3X-His and ubiquitin mutants restricted to K27, K48, or K63 linked ubiquitin chains and observed SYVN1 promoted K27- and K63-linked ubiquitination of DDX3X (Figure 4b). To determine whether distinct ubiquitin linkages influence cellular outcomes, HEK293T cells expressing TLR4 were transfected with SYVN1, DDX3X, and either K27- or K63-linked ubiquitin, then stimulated with LPS followed by nigericin or arsenite to induce inflammasome activation or stress granule formation, respectively. Nigericin treatment promoted both K27- and K63-linked ubiquitination of DDX3X by SYVN1, whereas arsenite stimulation selectively diminished K63 linkage without affecting K27 linked ubiquitination (Figure 4c). These results indicate stimulus-dependent remodeling of DDX3X ubiquitination, particularly the selective retention of K63-linked chains during inflammasome activation, may determine the functional availability of DDX3X for NLRP3 inflammasome assembly versus stress-granule assembly. We next sought to identify the lysine residues on DDX3X required for K63-linked ubiquitination. To predict the ubiquitinated region, we first examined ubiquitination of individual DDX3X domains to narrow down the modified area before residue-level mapping (Supplementary figure 4d). Di-glycine remnant profiling identified K554 as a prominent ubiquitination site (Supplementary figure 4e-f). However, mutation of K554 alone had no effect on total DDX3X ubiquitination (Figure 4e), suggesting that SYVN1 may target multiple functionally redundant lysine residues. Structural analysis of the AlphaFold model of DDX3X (UniProt ID: Q62167) revealed 32 lysine residues distributed across the protein surface. Our previous Di-glycine remnant profiling identified K554 as a prominent ubiquitination site and suggested that additional lysine residues located on the same protein surface might also contribute to ubiquitination. This surface contains eight additional lysine residues like K162, K208, K418, K491, K511, K581, K589 and K564 which are distributed across multiple domains but cluster spatially around K554 in the three-dimensional structure. To test whether this lysine-rich surface contributes to ubiquitination, we generated ubiquitination-deficient mutants by substituting lysine residues with arginine (K to R). Notably, the 9KR mutant exhibited markedly reduced K63-linked ubiquitination (Figure 4e). These results suggest that this spatially clustered lysine-rich surface preferentially supports K63-linked ubiquitination of DDX3X. To determine the functional significance of this modification, we reconstituted DDX3X-deficient BMDMs with wild-type DDX3X or mutant DDX3X (9KR). DDX3X wild type (WT) restored caspase-1 activation, GSDMD cleavage and IL-1β secretion whereas the mutant DDX3X (9KR) failed after treating the cells with LPS and nigericin to induce NLRP3 inflammasome activation (Figure 4f–g). However, both wild-type DDX3X and mutant DDX3X (9KR) restore stress granule assembly after arsenite stimulation (supplementary figure 4h-i). overall, these results demonstrate that K63-linked ubiquitination of DDX3X by SYVN1 is essential for NLRP3 inflammasome assembly and dispensable for stress granule assembly.

**Figure 4.**
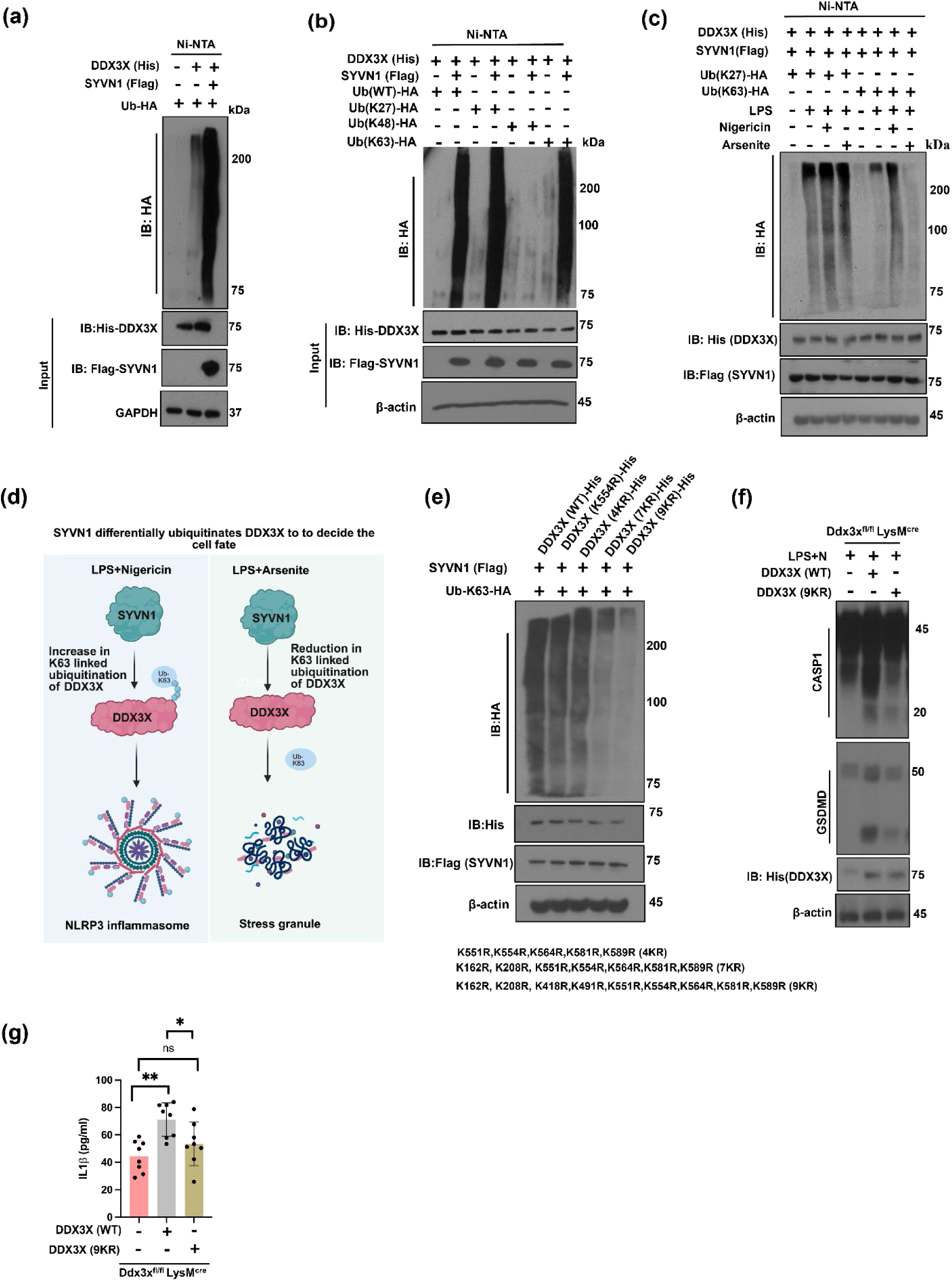
SYVN1 regulates NLRP3 inflammasome through K63 linked ubiquitination of DDX3X. (a) Immunoblot analysis of HEK293T cells transfected with plasmids for DDX3X-His, SYVN1-Flag and HA-tagged ubiquitin followed by immunoprecipitation with Ni-NTA beads (representative blots, n=3). (b) Immunoblot analysis of HEK293T cells transfected with plasmids for DDX3X-His, SYVN1-Flag and HA-tagged WT, K27, K48 and K63 mutant ubiquitin expressing plasmids followed by immunoprecipitation with Ni-NTA beads (representative blots, n=3). (c) Immunoblot analysis of HEK293T cells transfected with plasmids for DDX3X-His, SYVN1-Flag and HA-tagged K27 and K63 mutant ubiquitin expressing plasmids followed by treatment with 1μg/ml LPS and 20μM nigericin or 50μM arsenite and immunoprecipitation with Ni-NTA beads (representative blots, n=3). (d) Schematic showing K63 linked ubiquitination promotes NLRP3 inflammasome activation (e) Immunoblot analysis of HEK293T cells transfected with plasmids for DDX3X-His or Mutants DDX3X-His (K554R, 4KR, 7KR and 9KR) SYVN1-Flag and HA-tagged K63 ubiquitin followed by immunoprecipitation with Ni-NTA beads (representative blots, n=3). (f) Immunoblot analysis of CASP1 and GSDMD cleavage in DDX3X depleted cells transfected with wildtype and mutant DDX3X (representative blots, n=3). (g) Enzyme-linked immunosorbent assay measurement of IL-1β secreted from DDX3X depleted cells transfected with WT and K9R DDX3X (n=8). P values: *p<0.05, **p<0.01 and ns = not significant.

### OTUB1 Removes K63 Linkage to Promote Stress Granule Formation

We next sought to determine how SYVN1-dependent ubiquitination of DDX3X is differentially regulated to direct DDX3X toward either the NLRP3 inflammasome or stress granules. Because K63-linked ubiquitination of DDX3X was selectively reduced during stress granule formation, we reasoned that a stimulus-responsive deubiquitinase might mediate this switch. Based on prior proteomic reports, we screened OTUB1 as a deubiquitinase associated with stress granule formation [15]. co-immunoprecipitation revealed that the interaction between OTUB1 and DDX3X was markedly enhanced following arsenite treatment but diminished upon nigericin stimulation (Figure 5b). Depletion of OTUB1 restored K63-linked ubiquitination of DDX3X in arsenite-treated cells (Figure 5c). Consistentently, OTUB1 deficiency impaired stress granule formation (Figure 5d-e) but did not affect ASC speck formation and caspase1 cleavage (Supplementary figure 5a-c) indicating that OTUB1 selectively promotes the stress granule arm of DDX3X function. We next investigated how SYVN1 and OTUB1 coordinate with eachother in response to distinct stimuli. Co-immunoprecipitation revealed that SYVN1 interacted with OTUB1 following LPS stimulation and enhanced after nigericin treatment while the interaction diminished followed by arsenite stimulation (Supplemnetary figure 5e). Consistent with stimulus-dependent engagement of this complex, ubiquitination of OTUB1 was increased under nigericin stimulation but substantially reduced following arsenite treatment (Supplementary figure 5e). Together, these findings identify a stimulus-dependent SYVN1–OTUB1 regulatory axis that regulate differential ubiquitination of DDX3X to determine the cell fate.

**Figure 5.**
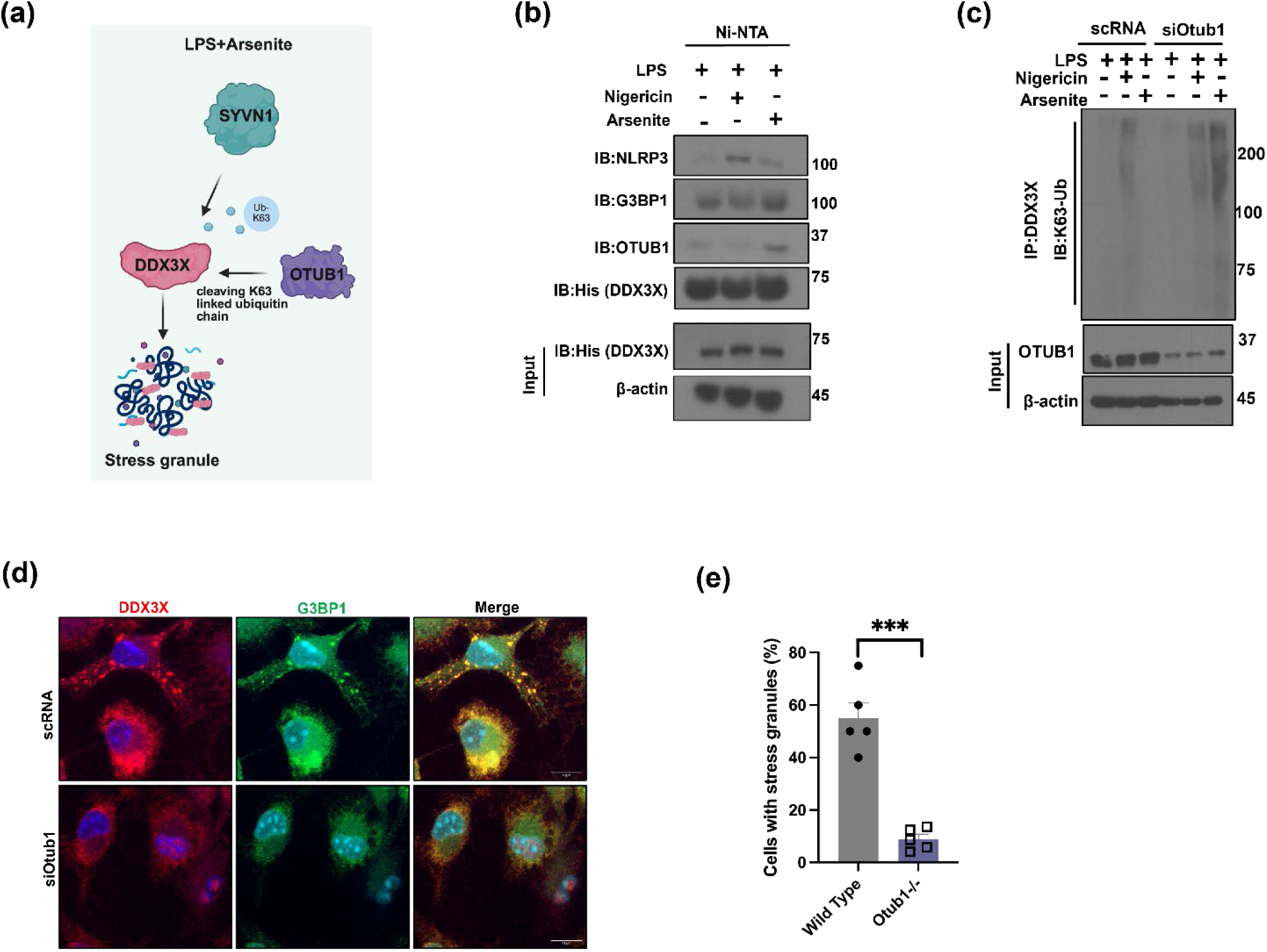
OTUB1 selectively cleaves K63 linked ubiquitin chain to facilitate stress granule assembly. (a) Schematic shows the OTUB1 cleaves K63 linked ubiquitin chain from DDX3X in macrophages stimulated with LPS and arsenite. (b) Immunoblot analysis of NLRP3, G3BP1 and OTUB1 in BMDM cells transfected with plasmid for DDX3X-His followed by treatment with 1μg/ml LPS with 20μM nigericin or 50μM arsenite and immunoprecipitation with Ni-NTA beads (representative blots, n=3).(c) Immunoblot analysis of K63 linked ubiquitination of DDX3X in siRNA depleted Otub1 compared with WT (representative blots, n=3).(d-e) Confocal images of stress granules in control and siRNA mediated depletion of Otub1 in BMDMs after treatment with LPS and arsenite (representative images n=5). percentage of stress granule positive cells stimulated with LPS and arsenite in wild type and OTUB1 depleted BMDMs. P value: ***p<0.001. Data are mean ± s.e.m.

## Discussion

The NLRP3 inflammasome and stress granules are cytosolic biomolecular condensates that enable cells to adapt to diverse forms of stress, promoting inflammatory cell death or cell survival respectively[1, 3, 4, 12, 13]. Recently, DDX3X has been shown as a shared component of these complexes and acts as a molecular checkpoint linking stress granule assembly to NLRP3 inflammasome activation [18]. However, it is unclear how DDX3X interpret the signal and choose between NLRP3 inflammasome mediated pyroptosis versus stress granule mediated cell survival. We speculated specific modifications on DDX3X drive the decision for cell fate. Studies showed E3 ligases play critical role in diverse cellular pathways essential for maintaining homeostasis. Several E3 ligases have been shown to regulate DDX3X activity either by proteasomal degradation or stabilizing the protein. TRIM36, RNF39, KLHL29, and WWP2 promote K48-linked ubiquitination and proteasomal degradation of DDX3X to tune macrophage polarization and RIG-I-like receptor (RLR) antiviral signaling [49–52], whereas TRIM25 promotes K63-linked ubiquitination of DDX3X to activate RLR signaling and antiviral defense [53]. In our study, we identified DDX3X as an interacting partner of SYVN1, so we investigated how SYVN1 influences DDX3X modification for selecting these contrasting cell fate decisions. SYVN1 is mainly associated with endoplasmic reticulum associated degradation (ERAD) pathway. However, SYVN1 acts as multifunctional protein and has an expanding functional repertoire in innate immunity, regulating T- and B-cell responses, their maturation and NF-KB–dependent inflammation through ubiquitination and inactivation of USP15 independent of ERAD [41]. More recently, SYVN1 was identified as an E3 ligase for GSDMD, mediating K27-linked ubiquitination on residues K203 and K204 to promote pyroptosis indicating the importance of SYVN1 in cell death pathway [42] and maintain SG homeostasis in stress condition [35]. The ability of SYVN1 to regulate both NLRP3 inflammasome activation and stress granule formation distinguishes it from previously described E3 ligases that regulate DDX3X in a more pathway-specific manner. Our study showed SYVN1 control Ddx3x mRNA expression either by degrading a repressor or activating the transcription factor to sustain the DDX3X transcription. We have shown that SYVN1 not only regulates the transcription of DDX3X, but it also ubiquitinates DDX3X through K63 linkage to determine the cell fate depending on the stimulus. Importantly, K63-linked ubiquitination of DDX3X is selectively required for the NLRP3 inflammasome activation, whereas DDX3X lacking this modification retains its ability to support stress granule formation. Although SYVN1 promoted both K27- and K63-linked ubiquitination of DDX3X. However, the presence or absence of K63-linked ubiquitination was functionally relevant for activating NLRP3 inflammasome or stress granule assembly. The identification of a spatially clustered lysine-rich surface encompassing K554 and eight additional lysine residues further suggests that DDX3X contains a structurally defined platform for K63-linked ubiquitination. Most importantly, reconstitution of DDX3X-deficient macrophages with the 9KR mutant failed to restore caspase-1 activation, GSDMD cleavage and IL-1β secretion, yet retained the ability to restore stress granule formation. These findings indicate that K63-linked ubiquitination of DDX3X become competent for inflammasome activation.

Our results also identify OTUB1 as a counter-regulatory component of this pathway. OTUB1 is an OTU-family deubiquitinase with established roles in ubiquitin signaling and innate immunity [48, 54, 55]. We found that OTUB1 associates more strongly with DDX3X under arsenite-induced stress and cleaved K63-linked ubiquitination of DDX3X. whereas this interaction is diminished following nigericin stimulation. Consistently, OTUB1 depletion restored K63-linked ubiquitination of DDX3X during arsenite treatment and selectively impaired stress granule formation without substantially affecting ASC speck formation or caspase-1 cleavage. These observations support a model in which OTUB1 promotes the stress granule-associated state of DDX3X by removing K63-linked ubiquitin chains. The stimulus-dependent relationship between SYVN1 and OTUB1 provides a potential mechanism for controlling this ubiquitin switch. Under inflammatory conditions, SYVN1 interaction with OTUB1 and SYVN1-dependent ubiquitination of OTUB1 were increased, whereas both were reduced following arsenite treatment. These reciprocal changes were accompanied by corresponding changes in the interaction of OTUB1 with DDX3X and in K63-linked ubiquitination of DDX3X. Thus, stimulus-dependent coordination between SYVN1 and OTUB1 establishes the ubiquitination state of DDX3X and provides a molecular switch that biases DDX3X toward NLRP3 inflammasome activation or stress granule assembly.

The physiological relevance of this regulatory pathway is supported by our in vivo findings. Myeloid-specific SYVN1 deficiency reduced IL-1β production and protected mice from LPS-induced acute lung injury, consistent with impaired NLRP3 inflammasome activation in vitro. At the same time, SYVN1 deficiency impaired stress granule formation under stress conditions, demonstrating that SYVN1 regulates both arms of the DDX3X-dependent stress response. These findings are particularly relevant to inflammatory diseases in which excessive NLRP3 inflammasome activation contributes to tissue injury, including sepsis and acute lung injury. More broadly, because dysregulated stress granule dynamics have also been implicated in chronic cellular stress and neurodegenerative disease.

### Limitations of the Study

Although our data established that K63-linked ubiquitination of DDX3X is required for NLRP3 inflammasome activation, the molecular mechanism by which this modification promotes DDX3X-dependent inflammasome assembly remains unclear. Future structural and biochemical studies will be required to determine whether K63 ubiquitination alters DDX3X conformation, protein–protein interactions, localization, or recruitment to the inflammasome. Addressing these questions will further define how stimulus-dependent ubiquitin remodeling of DDX3X controls the balance between inflammatory activation and stress adaptation.

While our study provides mechanistic insight into the SYVN1–DDX3X–OTUB1 axis, several limitations should be acknowledged. First, the role of OTUB1 was primarily investigated using siRNA-mediated depletion, and the absence of a myeloid-specific Otub1 conditional knockout model limits the physiological validation of this pathway in vivo. Future studies using Otub1^fl/fl^LysM^cre^ mice will be important to establish the in vivo contribution of OTUB1 to DDX3X ubiquitination and the balance between inflammasome activation and stress granule formation. Studies using catalytically inactive OTUB1 mutants, such as C91S, could help validating these mechanisms. SYVN1–OTUB1 association and OTUB1 ubiquitination are enhanced under inflammatory conditions and reduced following arsenite treatment; however SYVN1-mediated ubiquitination directly regulates OTUB1 catalytic activity has not been established. Mapping the relevant OTUB1 ubiquitination sites and testing site-specific mutants together with direct deubiquitinase activity assays will be necessary to understand the coordination between SYVN1 and OTUB1.

## Materials and Methods

### EXPERIMENTAL MODEL AND SUBJECT DETAILS

#### Cell Lines

Mouse bone marrow-derived macrophages (BMDMs) and HEK293T cells were used in this study.

#### Mice

C57BL/6 mice were obtained from Charles River Laboratories. Syvn1^fl/fl^ mice were purchased from Shanghai Model Organisms. Syvn1^fl/fl^LysM^cre^ mice were generated by crossing SYVN1^fl/fl^ mice with LysM^cre^ mice to achieve myeloid-specific deletion of SYVN1. Ddx3x^fl/fl^ and Ddx3x^fl/fl^LysM^cre^ mice were generously provided by Dr. Thirumala-Devi Kanneganti (St. Jude Children’s Research Hospital) [18]. All mice were housed in the University of Illinois Animal Care Facility in accordance with institutional and NIH guidelines. All animal experiments were approved by the University of Illinois Institutional Animal Care and Use Committee (IACUC). Male and female mice aged 6–8 weeks were randomly assigned to experimental groups.

### METHOD DETAILS

#### Cell Culture

BMDMs were differentiated from bone marrow progenitors over 6 days in DMEM (Gibco) supplemented with 10% fetal bovine serum (FBS; Gibco), 30% L929-conditioned medium as a source of M-CSF, and 1% penicillin-streptomycin (Sigma). Prior to stimulation, BMDMs were seeded overnight in antibiotic-free medium. HEK293T cells were maintained in DMEM supplemented with 10% FBS and 1% penicillin-streptomycin.

#### Inflammasome and Stress Granule Stimulation

To assess canonical NLRP3 inflammasome activation, BMDMs were primed for 3 h with 1 µg/ml ultrapure LPS (E. coli 0111: B4) (InvivoGen, tlrl-3eblps) and subsequently stimulated for 30–45 min with 20 µM nigericin (Cayman Chemical, 11437), 5 mM ATP (A2383, Sigma), or 10 µg/ml imiquimod (InvivoGen, tlrl-imqs-1). To induce stress granule formation, BMDMs were first primed with LPS (1 µg/ml) for 3 h and subsequently treated with 50 µM sodium arsenite (Sigma, S7400).

#### Immunoblot Analysis

Cells were washed with ice-cold PBS and lysed in RIPA buffer and samples were prepared in SDS containing sample-loading buffer with 2-mercaptoethanol. Proteins were resolved on SDS-polyacrylamide gels and transferred onto PVDF membranes. Membranes were blocked in 5% skimmed milk in TBST (0.1% Tween-20) for 1 h at room temperature, then incubated overnight at 4°C with the following primary antibodies: anti-SYVN1 (Proteintech, 67488-1-Ig), anti-caspase-1 p20 (AdipoGen, AG-20B-0042), anti-NLRP3 (AdipoGen, AG-20B-0014), anti-NLRP3 (Cell Signaling Technology, 15101), anti-ASC (AdipoGen, AG-25B-006-C100), anti-DDX3X (Santa Cruz Biotechnology, sc-365768), anti-G3BP1 (Proteintech, 66486-1-Ig), anti-GSDMD (Abcam, ab209845), anti-HA (Proteintech, 51064-2-AP), anti-FLAG (Sigma, F1804), anti-β-actin (Santa Cruz Biotechnology, sc-58673) and anti-OTUB1 (Proteintech, 68489-1-Ig). After washing with TBST (0.1%), membranes were incubated for 1 h at room temperature with the appropriate HRP-conjugated secondary antibody: anti-rabbit IgG (Jackson ImmunoResearch, 111-035-047), anti-mouse IgG (Jackson ImmunoResearch, 315-035-047). Proteins were detected using Luminata Forte Western HRP Substrate (Millipore, WBLUF0500).

#### Immunoprecipitation studies

For co-immunoprecipitation, cells were lysed in NP-40 lysis buffer (1% NP-40, 150 mM NaCl, 50 mM HEPES, pH 7.4) supplemented with protease and phosphatase inhibitor cocktails, and lysates were cleared by centrifugation at 12,000 rpm for 15 min at 4°C. Whole-cell lysates were incubated with Ni-NTA beads or 2 µg of the indicated primary antibody on a rocking platform overnight at 4°C. Pre-washed Protein A/G PLUS-Agarose beads (Santa Cruz Biotechnology) were blocked with 3% BSA for 30 min and then added to each sample for an additional 2 h at 4°C with rocking. Beads were collected by centrifugation and washed three to five times with TBST (0.1%). Bound proteins were eluted in SDS sample buffer and analyzed by immunoblotting.

To map the interaction interface between DDX3X and SYVN1, a panel of DDX3X constructs was used: full-length DDX3X-His, DDX3X-ΔN-term-His (residues 156–662), DDX3X-ΔC-term-His (residues 1–573), and DDX3X-helicase-His (residues 156–573). HEK293T cells were transiently co-transfected with 4 µg of the indicated DDX3X construct and 4 µg of Flag-tagged SYVN1 using Lipofectamine 2000 in Opti-MEM. Medium was replaced with DMEM containing 10% FBS 6 h post-transfection and cells were harvested 48 h post-transfection in NP-40 lysis buffer (1% NP-40, 150 mM NaCl, 50 mM Tris-HCl, pH 7.4, with protease and phosphatase inhibitors). Soluble lysates were subjected to immunoprecipitation with Ni-NTA beads. pCMV-3Tag-1A-P2A-SYVN1, pcDNA 3.1-myc-His-DDX3X and their domains were purchased from GenScript. OTUB1 expressing construct was purchased from Sino Biological (MG51912-NF). These constructs were used for coimmunoprecipitation studies.

#### In Vivo Ubiquitination Assay

To assess SYVN1-mediated ubiquitination of DDX3X, HEK293T cells were transfected with HA-tagged ubiquitin (wild-type, K27-only, K48-only, or K63-only), His-tagged wild-type or lysine-mutant DDX3X and SYVN1-Flag. To check the ubiquitination of domains of DDX3X, Domain constructs of DDX3X with SYVN1 (FL) were transfected with ubiquitin K63 in HEK293T. Similarly, to assess the domain of SYVN1 responsible for interaction and ubiquitination of DDX3X, constructs of SYVN1 with DDX3X-FL were transfected in HEK293T. At 48 h post-transfection, cells were lysed in denaturing lysis buffer (6 M guanidine-HCl, 0.1 M Na₂HPO₄/NaH₂PO₄, 10 mM imidazole, pH 8.0) and sonicated or NP40 lysis buffer followed by centrifugation to clarify the lysates. The supernatant was incubated with nickel-NTA agarose beads for 3 h at room temperature to pull down His-tagged DDX3X under denaturing conditions. Beads were washed extensively and bound proteins were eluted in SDS sample buffer for immunoblot analysis with anti-HA to detect ubiquitinated DDX3X.

#### Affinity Purification–Mass Spectrometry (AP-MS)

To identify SYVN1-interacting proteins and candidate substrates, HEK293T cells were transfected with Flag-tagged SYVN1 (Flag-SYVN1) and incubated for 48 h. Cells were then stimulated with 1 µg/ml LPS for 3 h and lysed in NP40 lysis buffer (20 mM Tris-HCl, pH 7.5, 100 mM KCl, 1 mM EDTA, 0.1% NP-40, 10% glycerol, and protease inhibitor cocktail). Clarified whole-cell lysates were incubated with anti-Flag antibody overnight at 4°C, followed by incubation with Protein A/G agarose beads for 2 h. Beads were washed five times with TBST (0.1%), and bound proteins were eluted in SDS loading buffer, resolved by SDS-PAGE, and the excised gel lane was submitted to the University of Illinois at Chicago proteomics core facility for mass spectrometry analysis.

#### ASC Oligomerization Assay

BMDMs from wild-type and SYVN1-deficient mice were lysed in NP-40 lysis buffer supplemented with 1 mM DSP (dithiobis(succinimidyl propionate)) crosslinker (CovaChem) to preserve protein complexes. Lysates were centrifuged at 10,000 rpm for 15–20 min to separate the insoluble (pellet) from the soluble fraction. The insoluble fraction was washed once with NP-40 lysis buffer and resuspended in the same buffer. Both fractions were mixed with 2X non-reducing SDS loading buffer (without β-mercaptoethanol) and analyzed by SDS-PAGE and immunoblotting with anti-ASC antibody to assess ASC oligomeric status.

#### Analytical Ultracentrifugation (AUC) and Glycerol Gradient Sedimentation

Control and SYVN1 depleted BMDMs were lysed in NP-40 lysis buffer containing 1 mM DSP crosslinker after stimulating with LPS (1μg/ml) and nigericin (20μM). A linear 15– 45% glycerol gradient in NP-40 lysis buffer was prepared by sequential layering of decreasing glycerol concentrations in a centrifuge tube. Concentrated lysate (50 µl) was carefully layered on top of the gradient, followed by 50 µl silicone oil to prevent evaporation. Tubes were centrifuged for 5 h at 55,000 rpm at 4°C in an Optima TLX (Beckman Coulter). Deceleration was performed without braking, and tubes were immediately placed on ice. Fractions of approximately 50 µl were collected sequentially from the top of the gradient by careful pipetting, with the bottom fraction yielding approximately 125 µl. A total of 17 fractions were collected into a 96-well plate and processed for SDS-PAGE and immunoblot analysis.

#### Immunofluorescence Microscopy

After stimulating the WT,SYVN1 and DDX3X depleted BMDMs with LPS (1μg/ml) with nigericin(20μM) or arsenite (50μM), cells were fixed for 15 min at room temperature with 4% paraformaldehyde (SantaCruz Biotechnology, sc-281692) and permeabilized with 0.1% Triton X-100 for 10 min. Cells were then incubated overnight at 4°C with the following primary antibodies: anti-ASC (Millipore, 04-147); 1:250), anti-DDX3X (Bethyl Laboratories, A300-474A; 1:100), anti-NLRP3 (CST, 15101; 1:100), rat anti-NLRP3 (R&D systems, MAB7578; 1:100), and anti-G3BP1 (Proteintech, 27299-1-AP; 1:250). Cells were washed and incubated with the appropriate Alexa 594 Fluor-conjugated secondary anti rabbit IgG (Life Technologies, A11036; 1:250), Alexa Fluor 647-conjugated anti mouse IgG (Life Technologies, A-21235;1:250), Alexa Fluor 488-conjugated anti rat IgG (Life Technologies, A-11006;1:250), Alexa Fluor 488-conjugated anti mouse IgG (Life Technologies, A-11001;1:250) for 1 h at room temperature. Nuclei were counterstained with Hoechst 33342 (Invitrogen, 62249). After slides preparation, Images were acquired on a Leica (LSM710 and 880) confocal microscope and processed using imageJ.

#### LPS-Induced Endotoxemia Model

To study the role of SYVN1 in survival and NLRP3-driven inflammatory injury, mice were given a single intraperitoneal injection of LPS (*Escherichia coli* serotype O111:B4; 2630, Sigma) at a dose of 15 mg/kg body weight. Serum and lung tissue were collected at the indicated time points for cytokine quantification and histological analysis.

#### Cycloheximide-chase assay

BMDMs from WT and SYVN1^fl/fl^ LysM^cre^ were treated with cycloheximide (100μg/ml) for indicated time points.

#### Quantitative RT-PCR

Total RNA was extracted from BMDMs using the RNeasy Micro Kit (Qiagen,74104) according to the manufacturer’s instructions. cDNA was synthesized from 500 ng total RNA using the High-Capacity cDNA Reverse Transcription Kit (Applied Biosystems). Quantitative PCR was performed using SYBR Green Master Mix (Applied Biosystems) real-time PCR system according to the manufacturer’s protocol. The following primer sequences were used for the PCR: *Syvn1*: forward, 5′-[ CCAACATCTCCTGGCTCTTCCA]-3′; reverse, 5′-[ CAGGATGCTGTGATAAGCGTGG]-3′; *Ddx3x*: forward, 5′-[ CTATGCCTCCAAAAGGTGTCCG]-3′; reverse, 5′-[ AGACCCAACTCTTCCTACAGCC]-3′ Gene expression was calculated using the comparative ΔΔC_T method and normalized to the endogenous reference gene *18S rRNA*.

#### siRNA-Mediated Knockdown of Syvn1 and Otub1

siRNA targeting mouse *Syvn1* (Dharmacon, L-041789-01-0005), Otub1 (Dharmacon, L2-046308-01-0005) and a non-targeting control siRNA pool (Dharmacon, D-001810-10-20) were obtained from Dharmacon. siRNAs were transfected into BMDMs using either the Amaxa Mouse Macrophage Nucleofector Kit (Lonza) or RNAiMax (Invitrogen) according to the manufacturers’ protocols. Knockdown efficiency was assessed by immunoblot analysis of SYVN1and OTUB1 proteins levels 48–72 h post-transfection.

#### ELISA

Concentrations of mouse IL-1β, IL-18 and TNF-α in cell culture supernatants. IL-1β in serum and lung tissue homogenates were measured using Quantikine sandwich ELISA kits (R&D Systems) according to the manufacturer’s instructions. IL-18 was quantified using an MBL ELISA kit. Absorbance was measured at 450 nm, and cytokine concentrations were calculated by linear regression against a standard curve using GraphPad Prism.

#### Lung injury evaluation in mice

Mice were administered EBA via retro-orbital injection 20% body weight ratio with a max volume of 100 µl using a 29G insulin syringe (150 µl max volume) after anesthesia. 30 minutes after EBA administration mice were sacrificed by cervical dislocation and the lungs were harvested. The lung injury level is calculated based on the methods described previously [56]. Followed by i.v. EBA (Millipore Sigma E2129, St. Louis, MO), the lung is perfused using cold PBS. Both lobes are harvested and weighed (wet weight) upon harvest. After measuring the wet weight, the lobe is incubated at 60°C for 16-48h in formamide for EB extraction. After the incubation, the formamide solution’s optical density (OD) is measured to estimate the quantity of EB released and recorded. The lung injury level is measured by;

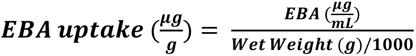

### QUANTIFICATION AND STATISTICAL ANALYSIS

All data are presented as mean ± SEM from at least three biologically independent experiments. Statistical comparisons were made using ordinary one-way ANOVA and 2-way ANOVA for multiple comparisons test in GraphPad Prism 11. Statistical significance between groups was indicated by asterisks in figures with the number of independent biological replicates indicated in each figure legend.

## ACKNOWLEDGMENTS

The studies were supported by NIH grants P01HL151327-03 (TO A.B.M). We thank Dr. Thirumala-Devi Kanneganti (St. Jude Children’s Research Hospital) for providing us Ddx3x^fl/fl^ and Ddx3x^fl/fl^LysM^cre^ mice strains.

## Author contribution

MA, A.B.M and C.T designed the studies; M.A, A.S, NRP and JT performed the experiments and the data analysis; and M.A, C.T and A.B.M. wrote the paper with critical input from the other authors.

---

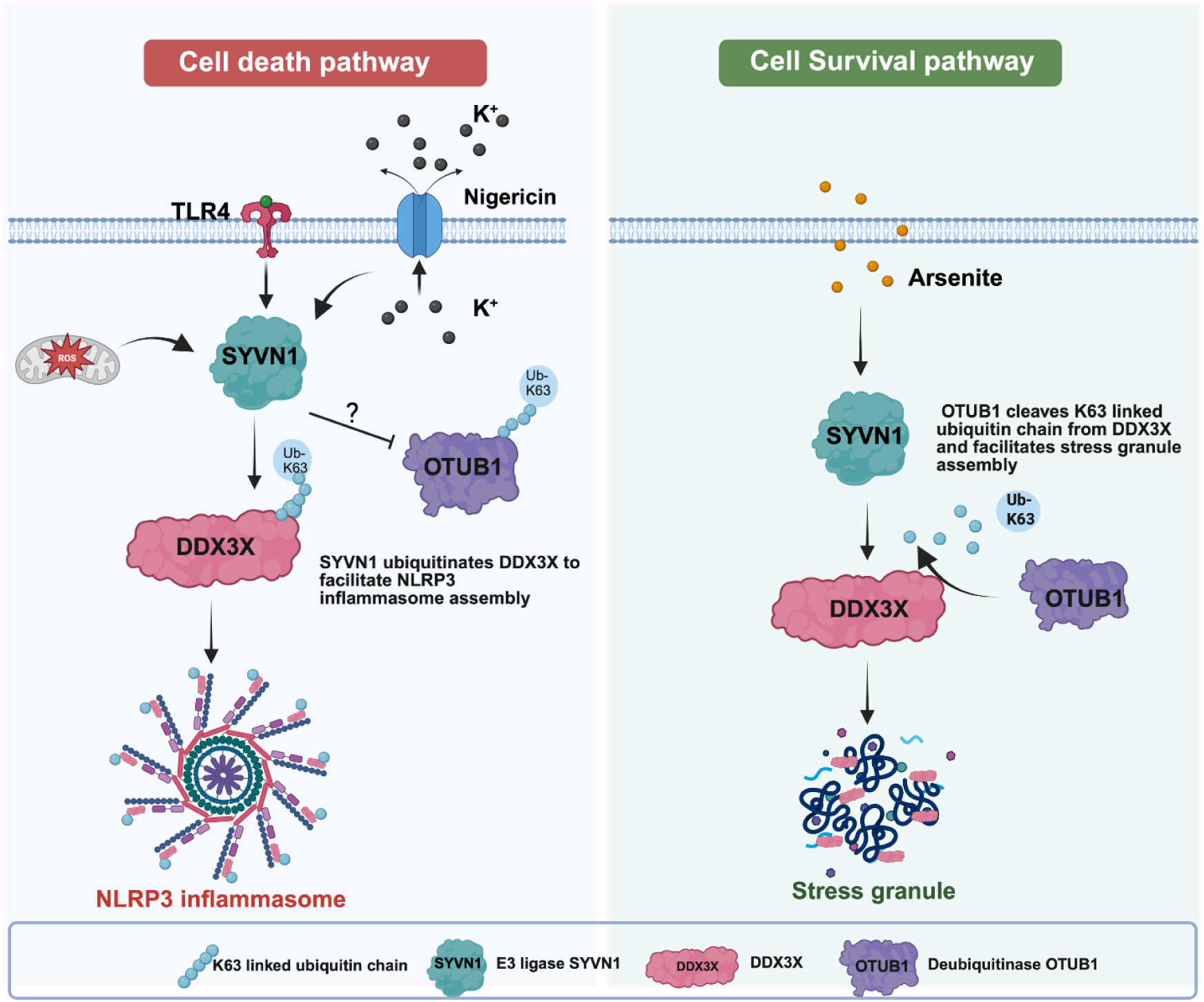
SYVN1-OTUB1 axis regulates K63-linked ubiquitination of DDX3X to switch cell fate between NLRP3 inflammasome activation and stress granule assembly. Schematic model shows the mechanism of NLRP3 inflammasome and stress granules assembly in response to cellular stress. DDX3X acts as a node and requires the activation of both condensates. During inflammation, DDX3X interacts with NLRP3 resulting in the activation of the inflammasome. Whereas, during other stress conditions, DDX3X facilitates the assembly of stress granules. However, the mechanism of how these condensates exploit the availibity of DDX3X is not fully understood. During inflammatory conditions, TLRs (Toll like receptors) recognize the signals and with K⁺ efflux or extreme oxidative condition resulting from excessive ROS production can stimulate the NLRP3 inflammasome activation. The activation of TLRs influences the expression of SYVN1 which controls the mRNA expression of Ddx3x and mediate K63 linked ubiquitination of DDX3X to regulate its function. In addition to that, SYVN1 interacts with deubiquitinase OTUB1 and prevents its interaction with DDX3X. Moreover, SYVN1 promotes OTUB1 ubiquitination as well that might lead to restricts its activity. The polyubiquitinated DDX3X interacts with NLRP3 resulting in the activation of inflammasome. Conversely, under arsenite-induced stress condition, the interaction between SYVN1 and OTUB1 diminished and thereby reduction in its ubiquitination resulting in enhanced interaction with DDX3X. OTUB1 cleaves K63 ubiquitin chain from DDX3X and favor cell survival by facilitating the stress

