## Supplementary Figures for "E3 ubiquitin ligase SYVN1 mediates K63-linked ubiquitination of DDX3X to activate Macrophage NLRP3 Inflammasome"

CORESSPONDENCE:

Asrar Malik,

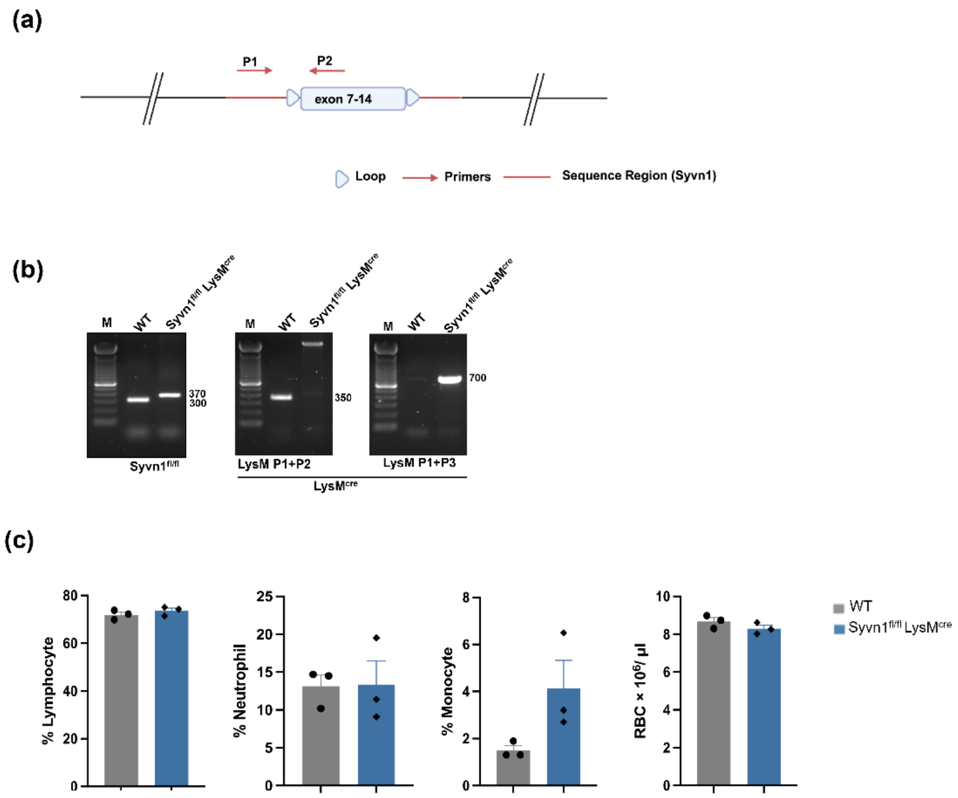


**Supplementary Figure 1.** (a) Schematic representation of genomic locations of loxP sites in the Syvn1 locus. (b) Genotyping PCR for the 5′ and 3′ loxP sites and LysM-cre. Representative gel (n = 1). (c) Quantification of the numbers of different immune-cell types from the blood of the indicated mouse strains. RBC, red blood cells (n=3). Data are mean ± s.e.m.

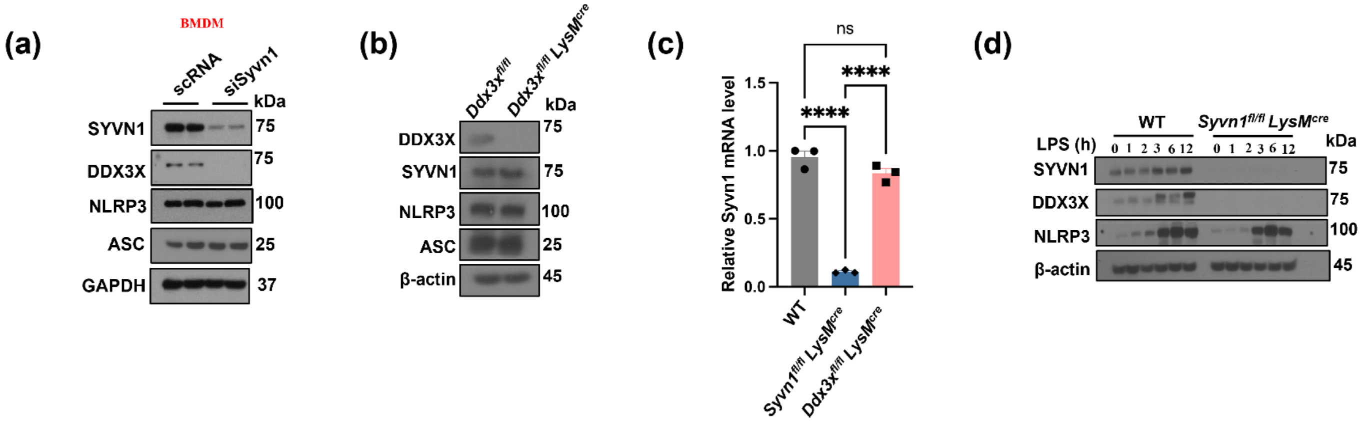


**Supplementary Figure 2. Unidirectional regulation of DDX3X expression by SYVN1.** (a) Immunoblot analysis of DDX3X, NLRP3, ASC protein expression in WT and SYVN1 depleted cells (representative images, n=3). (b) Immunoblot analysis of SYVN1, NLRP3, ASC protein expression in WT and DDX3X depleted cells (representative images, n=3). (c) qPCR analysis of the mRNA levels of Syvn1 in WT and Ddx3x$^{fl/fl}$LysM$^{cre}$ (n=3). P values: ****p< 0.0001 and ns = not significant. (d) Immunoblot analysis of SYVN1, DDX3X and NLRP3 protein expression in WT and SYVN1 depleted cells after treatment with LPS as indicated time points (representative images, n=3).

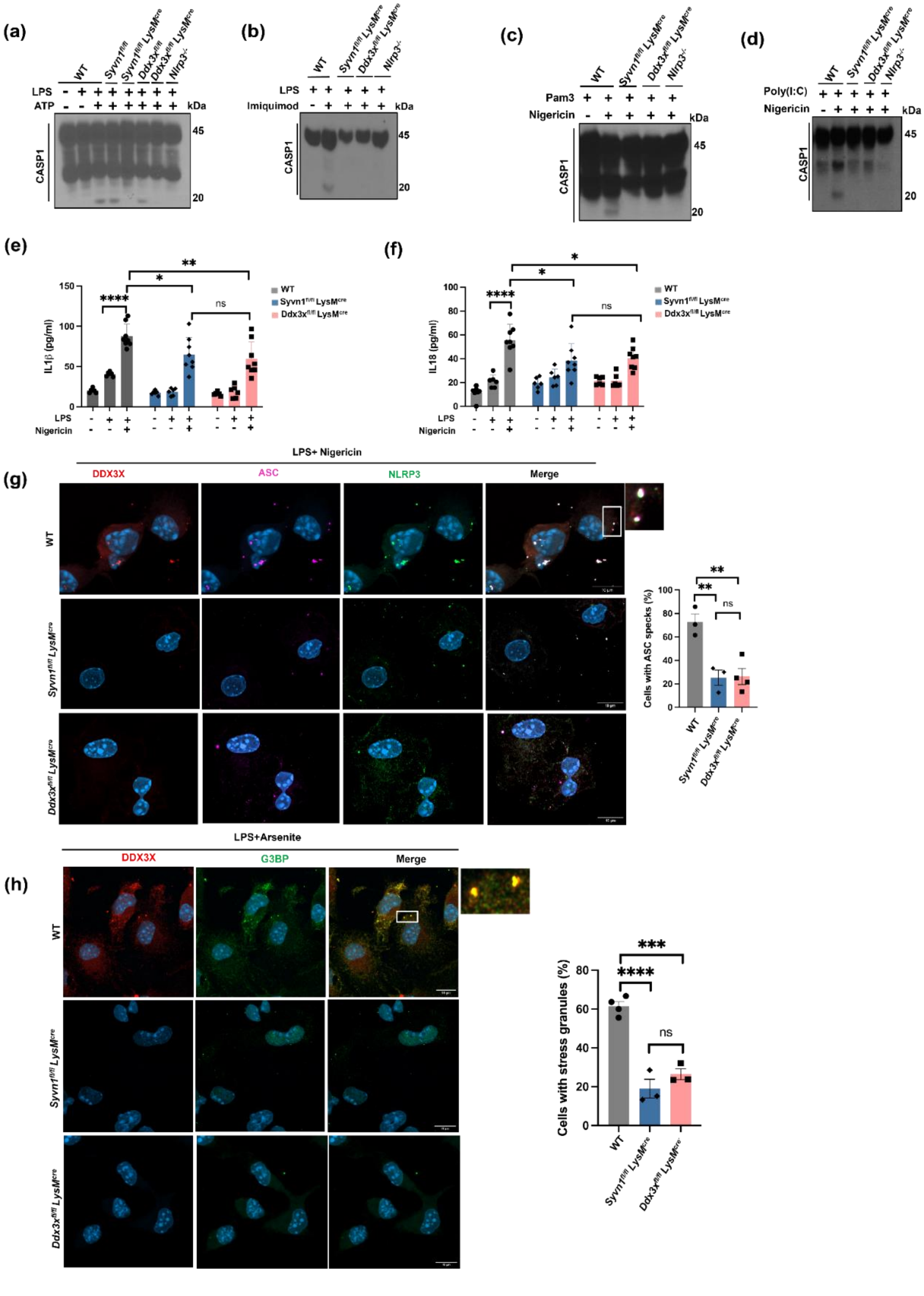


**Supplementary Figure 3. Lack of SYVN1 leads to defects in NLRP3 inflammasome activation and stress granule.** (a) Immunoblot analysis of CASP1 cleavage in control and siRNA mediated SYVN1 depleted BMDMs treated with LPS or LPS and ATP. (b) LPS or LPS and Imiquimod. (c) Pam3CSK4 or Pam3CSK4 and nigericin. (d) poly(I:C) or poly(I:C) and nigericin (representative blots, n = 2). (e-f) Enzyme-linked immunosorbent assay measurement of IL1β and IL18 from BMDMs isolated from WT, Syvn1$^{fl/fl}$ LysM$^{cre}$ and Ddx3x$^{fl/fl}$ LysM$^{cre}$ (n=7). P values: *p<0.05, **p<0.01 and ns = not significant. (g) Confocal images of NLRP3 inflammasome assembly in WT and compared with Syvn1$^{fl/fl}$ LysM$^{cre}$ and Ddx3x$^{fl/fl}$ LysM$^{cre}$. Scale bars,10μm (whole cell images); 1μm (magnified images) (representative images, n=3). Quantification of ASC speck positive cells stimulated with LPS and nigericin as indicated. P values **p< 0.01 and ns = not significant. Data are mean± s.e.m. (h) Confocal images of stress granules assembly in WT and compared with Syvn1$^{fl/fl}$ LysM$^{cre}$ and Ddx3x$^{fl/fl}$ LysM$^{cre}$. Scale bars,10μm (whole cell images); 1μm (magnified images) (representative images, n=4). Quantification of stress granule positive cells stimulated with LPS and arsenite as indicated. P values, ***p<0.001 and ****p<0.0001 and ns = not significant. Data are mean± s.e.m.

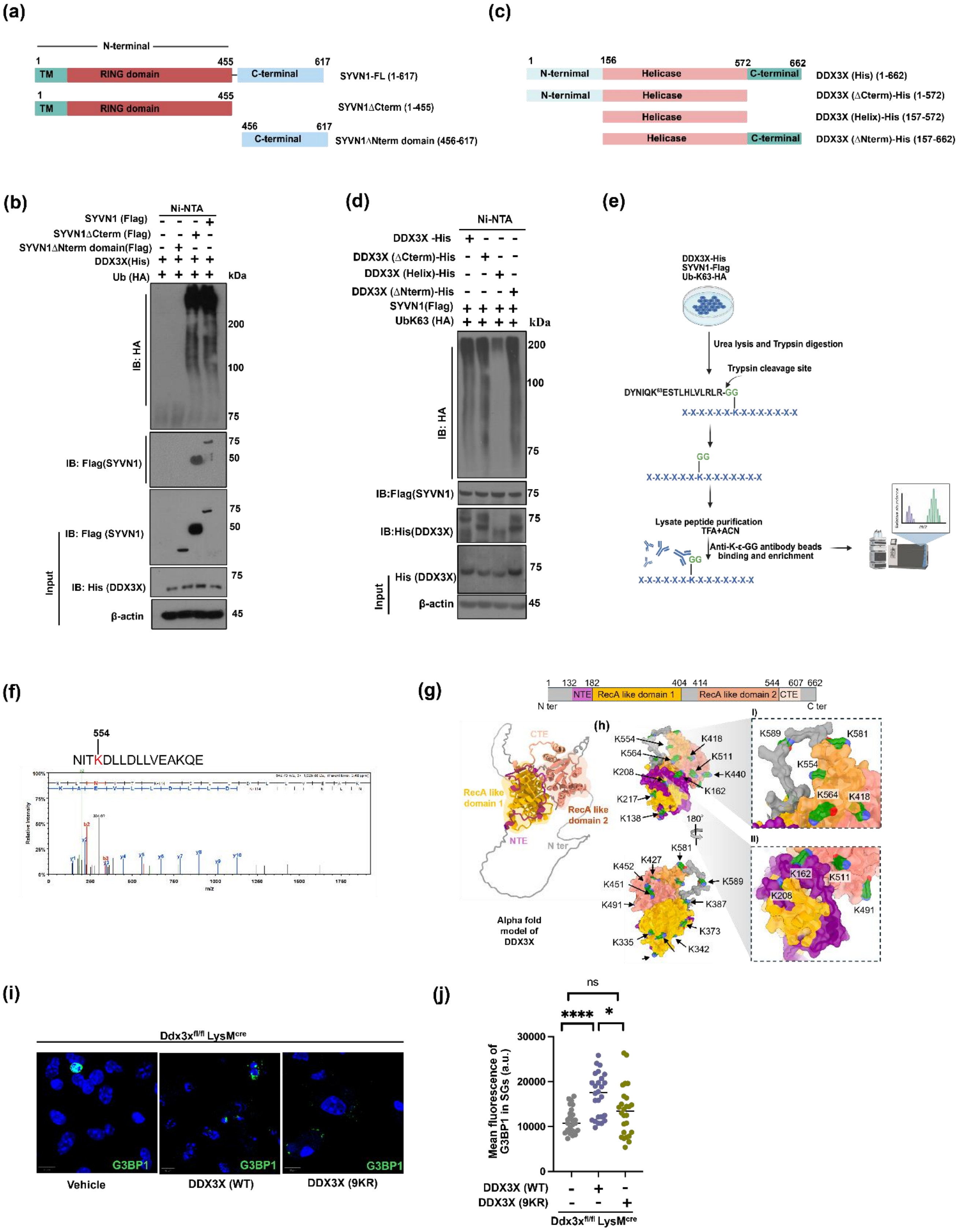
(a)
N-terminal
1
455
617
TM
RING domain
C-terminal
SYVN1-FL (1-617)
SYVN1ΔCterm (1-455)
456
SYVN1ΔNterm domain (456-617)
(c)
1
156
572
662
N-terminal
Helicase
C-terminal
DDX3X (His) (1-662)
DDX3X (ΔCterm)-His (1-572)
DDX3X (Helix)-His (157-572)
DDX3X (ΔNterm)-His (157-662)
(b)
Ni-NTA
SYVN1 (Flag)
SYVN1ΔCterm (Flag)
SYVN1ΔNterm domain(Flag)
DDX3X(His)
Ub (HA)
kDa
200
100
75
50
45
IB: HA
IB: Flag(SYVN1)
IB: Flag (SYVN1)
IB: His (DDX3X)
β-actin
Input
(d)
Ni-NTA
DDX3X -His
DDX3X (ΔCterm)-His
DDX3X (Helix)-His
DDX3X (ΔNterm)-His
SYVN1(Flag)
UbK63 (HA)
kDa
IB: HA
IB:Flag(SYVN1)
IB:His(DDX3X)
His (DDX3X)
β-actin
Input
(e)
DDX3X-His
SYVN1-Flag
Ub-K63-HA
Urea lysis and Trypsin digestion
Trypsin cleavage site
DYNIQK63ESTLHLVLRLR-GG
X-X-X-X-X-K-X-X-X-X-X-X-X
GG
Lysate peptide purification
TFA+ACN
Anti-K-ε-GG antibody beads binding and enrichment
(f)
554
NITKDLLDLLVEAKQE
(g)
1 132 182 404 414 544 607 662
NTE
RecA like domain 1
RecA like domain 2
CTE
N ter
C ter
RecA like domain 1
RecA like domain 2
Alpha fold model of DDX3X
(h)
K554
K564
K418
K511
K440
K208
K162
K217
K138
180°
K581
K452
K427
K589
K451
K387
K491
K373
K335
K342
I)
K589
K581
K554
K564
K418
II)
K162
K511
K208
K491
(i)
Ddx3xfl/fl LysMcre
G3BP1
Vehicle
DDX3X (WT)
DDX3X (9KR)
(j)
ns
****
*
Mean fluorescence of G3BP1 in SGs (a.u.)
30000
20000
10000
0
DDX3X (WT)
DDX3X (9KR)
Ddx3xfl/fl LysMcre

**Supplementary Figure 4. Domain mapping and identification of SYVN1-mediated K63-linked ubiquitination sites on DDX3X and their role in stress granule formation.** (a) Schematic representation of domain organization of SYVN1 constructs: full-length SYVN1 (SYVN1-FL), the N-terminal domain of SYVN1 containing transmembrane domain (TM) and Ring domain (SYVN1 ΔC terminal) and C-terminal domain of SYVN1 (SYVN1 ΔN terminal). (b) Immunoblot analysis of immunoprecipitation and ubiquitination of DDX3X by indicated SYVN1 domains (representative images, n=2). (c) Schematic representation of DDX3X domains as indicated. (d) Identification of the region of DDX3X required for its interaction with SYVN1 and their ubiquitination by SYVN1 (representative blots, n=2). (e) Identification of ubiquitination site on DDX3X: schematic representation of workflow for LC-MS analysis. (f) ubiquitinated "K" containing peptides of DDX3X identified by LC-MS. (g) Schematic representation of DDX3X showing its domain organization and AlphaFold-predicted structural model of DDX3X, with individual domains colored according to the domain organization. (h) Surface representation of DDX3X from different orientations, highlighting surface-exposed lysine residues in green. (h- I–II) enlarged views of the region surrounding K554, showing additional lysine residues located in spatial proximity to K554 and distributed across multiple domains of DDX3X. (i) Confocal images of stress granules assembly in BMDMs isolated from Ddx3x$^{fl/fl}$ LysM$^{cre}$ mice and transfected with wild type DDX3X and mutant DDX3X (9KR). Scale bars,10μm. (j) Fluorescence intensities of G3BP1 in stress granules (n=19). P values: *p<0.05, **p<0.01 and ns = not significant.

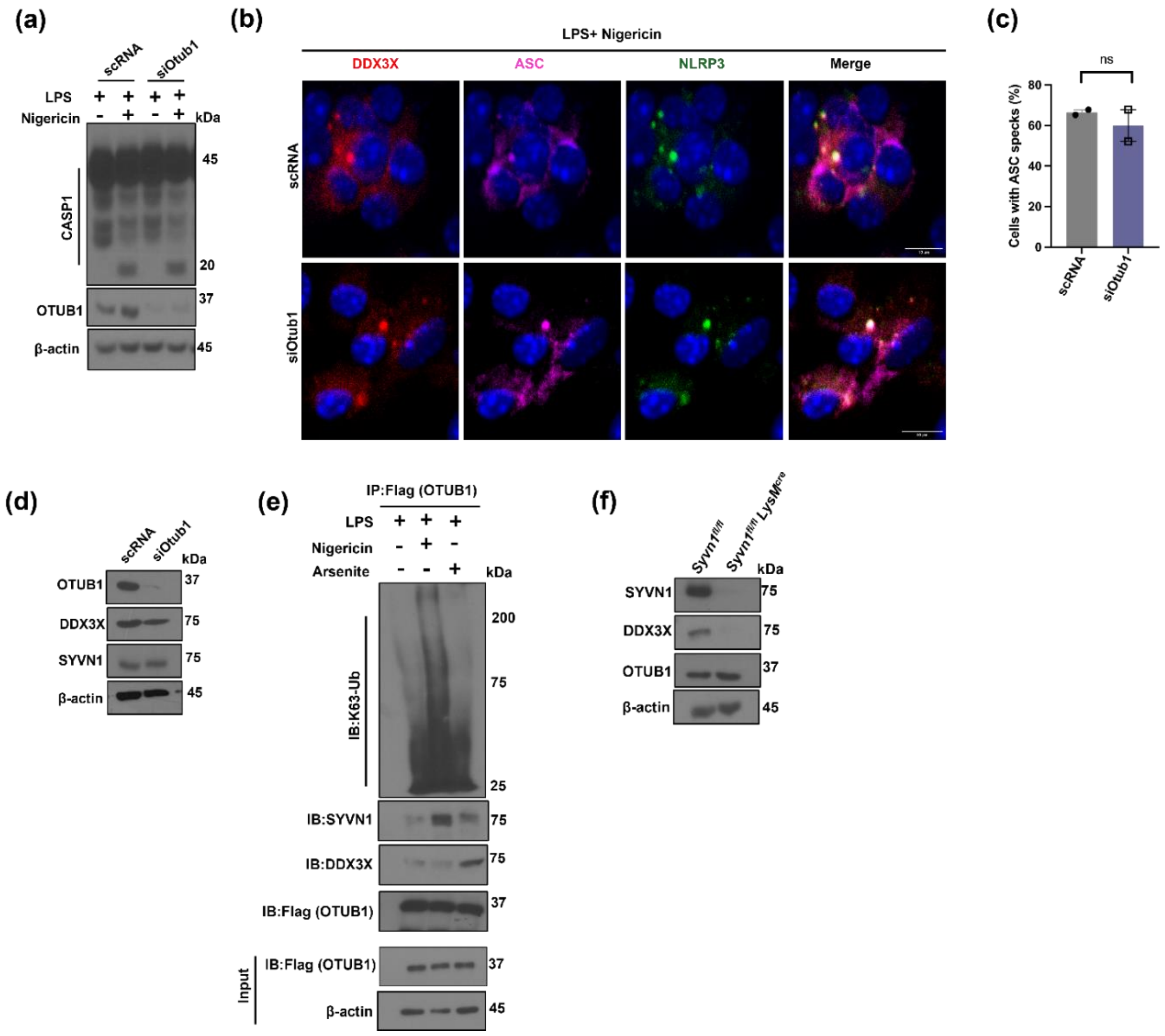


**Supplementary Figure 5. OTUB1 depletion had no effect on NLRP3 inflammasome activation.** (a) Immunoblot analysis of DDX3X and SYVN1 in control and siRNA mediated OTUB1 depleted BMDM cells (representative images, n=3). (b) Immunoblot analysis of CASP1 cleavage in control and siRNA mediated OTUB1 depleted BMDMs treated with LPS or LPS and Nigericin (c-d) Confocal images of NLRP3 inflammasome assembly in WT and compared with OTUB1 depleted cells. Scale bars,10μm (whole cell images) (represenative images, n=2). Quantification of ASC speck positive cells stimulated with LPS and nigericin as indicated. P value: ns = not significant. Data are mean ± s.e.m. (e) Immunoblot analysis of immunoprecipitation of SYVN1 and DDX3X and ubiquination of OTUB1 in cells stimulated with LPS, LPS and Nigericin or arsenite (representative image, n=3). Immunoblot analysis of DDX3X and OTUB1 in SYVN1 depleted cells (representative images, n=3).
